# runBioSimulations: an extensible web application that simulates a wide range of computational modeling frameworks, algorithms, and formats

**DOI:** 10.1101/2021.03.05.433787

**Authors:** Bilal Shaikh, Gnaneswara Marupilla, Mike Wilson, Michael L. Blinov, Ion I. Moraru, Jonathan R. Karr

**Affiliations:** Icahn Institute for Data Science and Department of Genetics and Genomic Sciences, Icahn School of Medicine at Mount Sinai, 1255 5th Avenue, Suite C2, New York, NY 10029, USA; Center for Cell Analysis and Modeling, University of Connecticut School of Medicine, 263 Farmington Avenue, Farmington, CT 06030, USA

## Abstract

Comprehensive, predictive computational models have significant potential for science, bioengineering, and medicine. One promising way to achieve more predictive models is to combine submodels of multiple subsystems. To capture the multiple scales of biology, these submodels will likely require multiple modeling frameworks and simulation algorithms. Several community resources are already available for working with many of these frameworks and algorithms. However, the variety and sheer number of these resources make it challenging to find and use appropriate tools for each model, especially for novice modelers and experimentalists. To make these resources easier to use, we developed runBioSimulations (https://run.biosimulations.org), a single web application for executing a broad range of models. runBioSimulations leverages community resources, including BioSimulators, a new open registry of simulation tools. These resources currently enable runBioSimulations to execute nine frameworks and 44 algorithms, and they make runBioSimulations extensible to additional frameworks and algorithms. runBioSimulations also provides features for sharing simulations and interactively visualizing their results. We anticipate that runBioSimulations will foster reproducibility, stimulate collaboration, and ultimately facilitate the creation of more predictive models.

## INTRODUCTION

More comprehensive and predictive models have significant potential for biology, bioengineering, and medicine. For example, models of entire cells could help synthetic biologists design cells and help physicians personalize medicine (1, 2).

Due to the complexity of biology, building such models will likely require collaboration among many modelers and experimentalists. One promising way to harness this diverse expertise is to combine multiple submodels of individual subsystems, each developed by a small group of experts.

To capture the multiple scales relevant to biology at a practical computational cost, these submodels will likely require multiple modeling frameworks. For example, a model of the phenotypic heterogeneity of single cells might need to capture slow processes, such as transcription, precisely, while fast processes, such as metabolism, could be captured coarsely. Numerous frameworks and simulation algorithms already exist for several scales. For example, stochastic kinetic simulations can capture the cell-to-cell variability in gene expression, and flux-balance analysis (FBA) can capture the distribution of fluxes over metabolic networks.

To facilitate collaboration, several model formats have been created for these frameworks and algorithms. For example, the BioNetGen Language (BNGL,3), CellML (4), NeuroML (5), and the Systems Biology Markup Language (SBML,6) can capture kinetic models, and the SBML flux balance constraints (SBML-fbc;7) and qualitative modeling (SBML-qual;8) packages can capture flux balance and logical models.

Furthermore, numerous software tools support these formats. For example, BioNetGen (9), NFSim (10), and Virtual Cell (VCell;11) support BNGL; BoolNet (12) and The Cell Collective (13) support SBML-qual; CBMPy (14), COBRApy (15), and The Cell Collective support SBML-fbc; COPASI (16), JWS Online (17), StochSS (18), tellurium (19), and VCell (11) support SBML; OpenCOR supports CellML (20); and Open Source Brain supports NeuroML (21).

These resources provide experts rich silos for modeling individual subsystems. However, this siloing poses obstacles to composing models of multiple subsystems and scales into comprehensive models. The effort required to learn the unique frameworks, algorithms, formats, software tools, and conventions associated with each silo also impedes collaboration, especially for novice modelers and experimentalists.

To help investigators use these resources, we developed runBioSimulations, an extensible REST API and graphical web application for executing simulations involving a broad range of frameworks, algorithms, and model formats. runBioSimulations leverages several community resources, including model formats such as SBML, the Simulation Experiment Description Language (SED-ML,22), the COMBINE archive format (23), and BioSimulators (https://biosimulators.org), a new open registry of standardized simulation tools. As of this writing, runBioSimulations supports nine frameworks, 44 algorithms, and five model formats (Table S1). This includes 37 algorithms for continuous, discrete, hybrid, and rule-based kinetic simulation with BNGL and SBML; five algorithms for flux balance simulation with SBML-fbc; and three algorithms for logical simulation with SBML-qual. Importantly, the community can expand runBioSimulations to additional frameworks, algorithms, and formats by contributing additional simulation tools to BioSimulators. runBioSimulations also provides features for visualizing simulation results and debugging, managing, and sharing simulations. Furthermore, the runBioSimulations API enables the community to develop additional front-end applications that utilize runBioSimulations’ unique simulation capabilities. For example, model repositories could use the API to provide interactive simulations of their models.

By making it easier to execute a broad range of models, we anticipate that runBioSimulations will foster model reuse, bolster collaboration, and empower peer review. In turn, we anticipate this will accelerate the development of more comprehensive and more predictive models.

Below, we describe the key features of runBioSimulations, its architecture, and how it facilitates model reuse and collaboration. In addition, we outline our future plans for runBioSimulations. The Supplementary Data summarizes the frameworks, algorithms, formats, and simulation tools supported by runBioSimulations; provides additional information about the implementation of runBioSimulations; presents a case study of using runBioSimulations to evaluate the practical reusability of existing published simulations to individual investigators that illustrates the utility of runBioSimulations; compares runBioSimulations to other tools; and outlines how the community can contribute to runBioSimulations.

## KEY FEATURES

The key feature of runBioSimulations is the capability to execute a broad range of simulations that involve a variety of modeling frameworks, simulation algorithms, and model formats from a single, simple, consistent interface. This is achieved through a modular architecture that leverages existing resources including model formats such as SBML, SED-ML, and the COMBINE archive format to encapsulate the details of each framework, algorithm, and model format. This architecture is implemented as a REST API. The runBioSimulations graphical user interface (GUI) provides investigators a user-friendly client to this powerful API.

The starting point to using runBioSimulations is a COMBINE archive that contains one or more models in a format such as BNGL or SBML and describes one or more simulations of these models in SED-ML. Users can obtain models and simulations encoded in these formats from repositories such as BioModels (24) or use tools such as VCell to create models and simulations in these formats. Models and simulations can be packaged into COMBINE archives using tools such as CombineArchiveWeb (25).

The runBioSimulations GUI enables users to execute models and retrieve and visualize their results in three simple steps. First, users use the GUI to select a COMBINE archive to execute and a simulation tool to run the archive (Figure2b). Users can choose any of the standardized simulation tools available in the BioSimulators registry and any of their versions. To help investigators find tools that are compatible with specific types of models and/or that support specific simulation algorithms, BioSimulators provides detailed information about the capabilities of each simulation tool (Figure 2a). The ability to use multiple simulators has several benefits. (a) This makes it easier to reuse models, including older models that require legacy formats. (b) This design makes the simulation logic of runBioSimulations transparent and portable, ensuring users that they can continue work initiated with runBioSimulations onto their own computers using the same simulators, further lowering the barrier to model reuse. (c) Because BioSimulators is an open registry, this design enables the community to extend the simulation capabilities of runBioSimulations.

**Figure 1.**
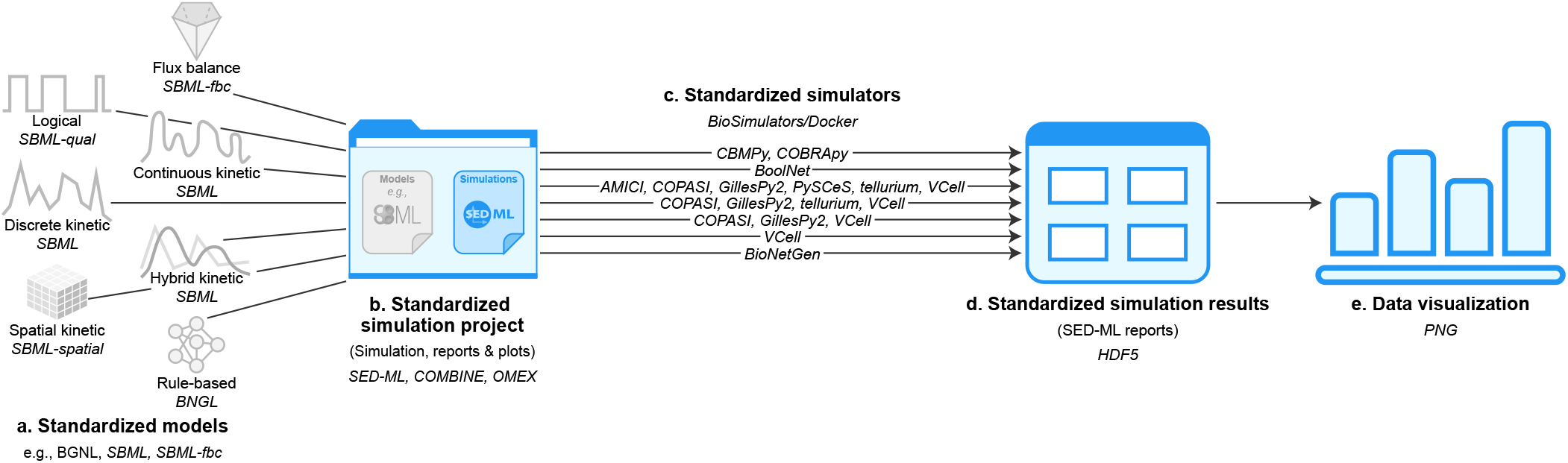
runBioSimulations is an extensible, standards-driven web application for executing models across a broad range of modeling frameworks, simulation algorithms, and model formats. runBioSimulations can execute models (**a**) and simulations (**b**) described with community resources such as COMBINE, SBML, and SED-ML with standardized simulation tools registered with BioSimulators (**c**). runBioSimulations produces results in HDF5 format (**d**). runBioSimulations also provides tools for interactively visualizing simulation results (**e**).

**Figure 2.**
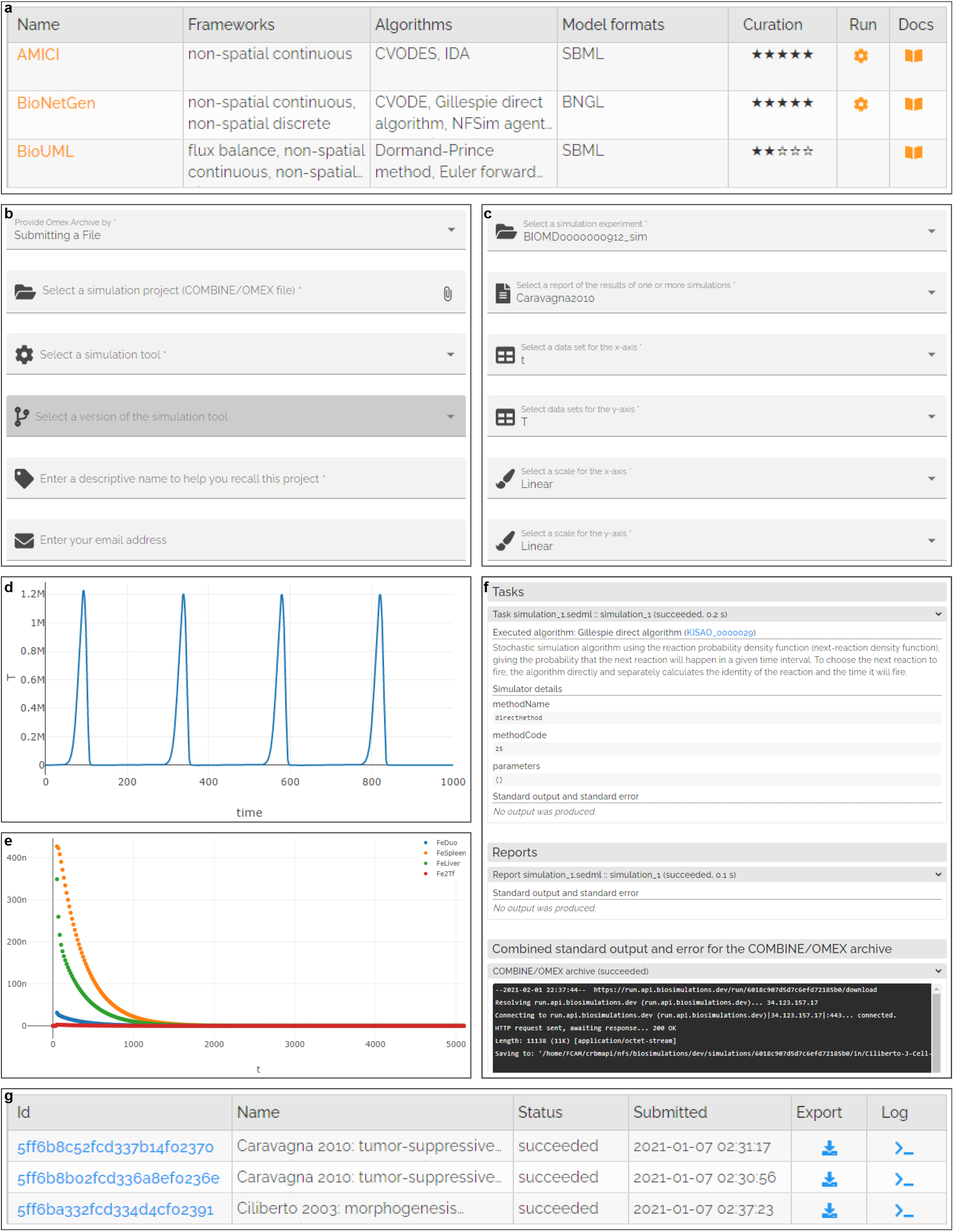
runBioSimulations provides a single GUI for executing a broad range of models and visualizing their results. (**a**) Users can use BioSimulators to select tools for executing specific simulations. (**b**–**e**) runBioSimulations provides simple forms for executing simulations and designing visualizations of their results. runBioSimulations also provides a summary table (**g**) and structured logs (**f**) for managing and debugging simulations.

Users can manage their simulations and monitor their progress using a table that summarizes their simulations (Figure 2g). Optionally, users can also provide an address to receive emails about the completion of the execution of their archives. This feature is valuable for long simulations.

Once simulations complete, users can download and visualize their results. Simulation results can be downloaded in HDF5 format. The GUI provides users a simple form (Figure 2c) for designing two-dimensional plots of model predictions (Figure 2d, e).

Users also have the option to upload Vega visualizations (26) to visualize simulation results. This enables investigators to visualize their simulation results with a broad range of charts, as well as custom, interactive, publication-quality diagrams. This also makes it easier to reuse visualizations across multiple simulation conditions by re-painting them with results of alternative simulations. Together, this combination of runBioSimulations and Vega ensures that the provenance of simulation results and visualizations of simulation results are transparent by capturing all of the information needed to reproduce each result and visualization, including the model, simulation, and simulator which generated each result and the transformations used to map each result to each diagram.

To help users debug simulations, the GUI also displays structured logs of their execution (Figure 2f). This can help direct users to errors in specific SED-ML tasks and outputs.

runBioSimulations also makes it easy for users to share simulations and their results via persistent URLs similar to file sharing services such as Google Drive. These links enable users to revisit their simulation results, share simulations with collaborators, anonymously share simulations with peer reviewers, and publish simulations by embedding links into articles. runBioSimulations is particularly well-suited to sharing computationally-expensive simulations because it enables investigators to quickly retrieve their results without having to wait for long simulations to complete.

Furthermore, developers can use runBioSimulations’ REST API to build additional client applications that leverage runBioSimulations’ simulation logic. For example, developers could use the API to build additional clients for executing simulations such as Jupyter notebooks or desktop applications.

## METHODS

runBioSimulations is composed of a GUI for submitting simulations, managing simulations, and visualizing their results; services for executing simulations on a high-performance computing (HPC) cluster, monitoring their progress, and collecting their results; and a database for storing simulations and their results (Figure 3). More information about the design, implementation, and deployment of runBioSimulations is available in the Supplementary Data.

**Figure 3.**
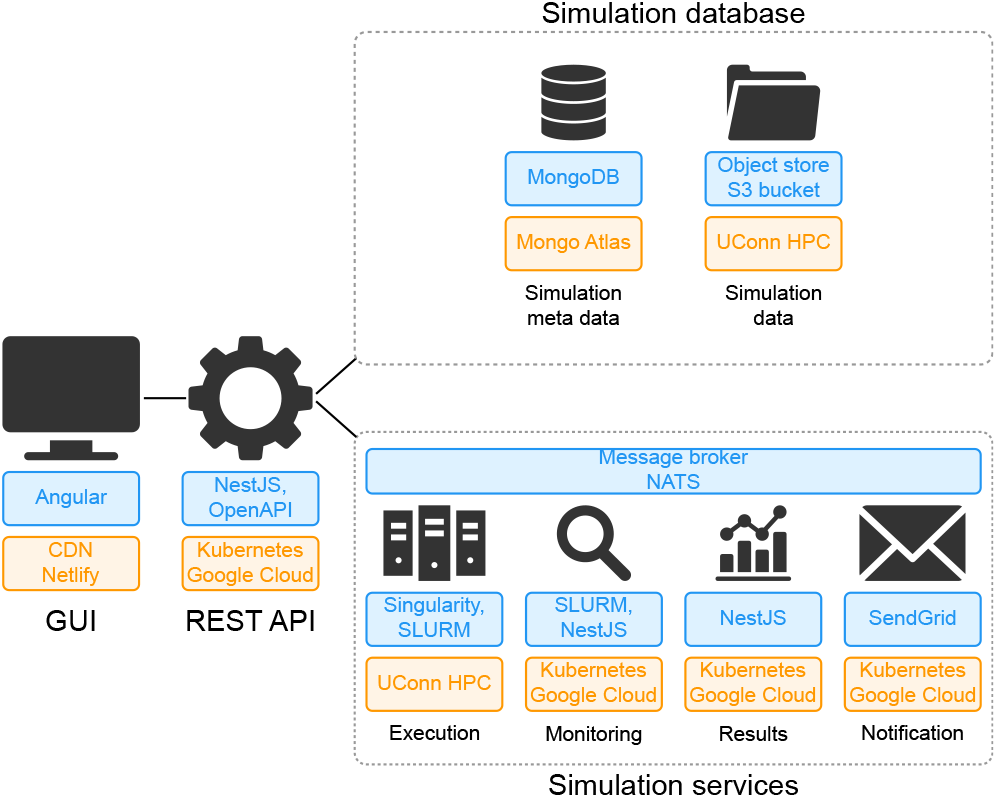
Overview of the implementation (blue) and deployment (orange) of runBioSimulations. The application consists of a GUI; services for executing, monitoring, and logging simulations; and a database of simulations and their results which interact via a REST API. To support multiple simultaneous users, runBioSimulations is deployed using the cloud and HPC.

## USE CASES

### Publishing simulations

We believe that runBioSimulations’ standards-driven design and unique capability to execute a broad range of simulations is ideal for publishing simulations. As more tools embrace SED-ML, runBioSimulations will help authors publish simulations that other investigators can easily reuse. For example, investigators could use runBioSimulations to explore additional conditions and predictions of a model beyond those reported by its authors. Furthermore, by providing simple access to multiple simulators, runBioSimulations can help authors and model curators verify that simulation results are reproducible across simulation algorithms and simulators.

### Collaboration and peer review

We believe that runBioSimulations is similarly well-suited for sharing simulations with collaborators and reviewers. By helping investigators work with different frameworks and algorithms, runBioSimulations makes it easier for investigators to contribute to multiple collaborations. In particular, runBioSimulations’ simulation URLs make it easy for investigators to share simulations with collaborators, and they enable peer reviewers to access simulations anonymously.

### Comparing simulation tools

Because runBioSimulations can execute the same simulations with multiple simulation tools, runBioSimulations is also well-suited to assessing the compatibility between tools. For example, investigators could compare results of the same simulations generated with multiple tools to evaluate the performance of the tools, identify inconsistencies among the tools, or detect potential errors in the tools.

### Multiscale modeling education

Furthermore, we believe that runBioSimulations could be a valuable educational tool. In particular, instructors who use runBioSimulations for assignments involving multiple frameworks would only need to teach their students a single tool. Instructors could also leverage runBioSimulations’ simulation results storage for assignments involving the analysis of results of computationally-expensive simulations.

## DISCUSSION

In summary, runBioSimulations provides a simple GUI for executing a broad range of simulations described using community resources such as SBML and SED-ML. Importantly, the community can extend these simulation capabilities by contributing additional standardized simulation tools to the BioSimulators registry. The runBioSimulations GUI also provides users features for managing their simulations, interactively visualizing their results, and sharing their simulations through persistent URLs. In addition, developers can use runBioSimulations’ API to build custom applications for executing simulations and/or analyzing simulation results. Together, we believe runBioSimulations will both help authors of in silico experiments share their simulations and help other investigators reproduce and reuse their studies. Ultimately, we believe runBioSimulations will facilitate collaboration and foster more comprehensive and more predictive models.

### Additional modeling formalisms, algorithms, and formats

We invite developers to extend runBioSimulations to more simulations by contributing additional standardized simulation tools to BioSimulators. To help developers standardize their tools, BioSimulators provides a Python library for executing COMBINE archives, a test suite for validating simulation tools, several examples, and documentation.

### More sophisticated data visualizations

We also aim to expand the visualization features of runBioSimulations by using Vega to support a broad range of canonical chart types, as well as custom charts, such as network maps. By capturing how charts can be painted with data, Vega would also enable users to reuse diagrams with multiple models and simulations, furthering our goals of reuse and collaboration.

### Online platform for sharing entire simulation projects

Furthermore, we plan to use the runBioSimulations API to develop an online platform that will help authors create and publish entire simulation studies and provide the community a central place to discover and reuse studies. This platform will layer several additional capabilities on top of runBioSimulations. The platform will enable authors to publish models, simulations, simulation results, and data visualizations of simulation results. The platform will also help the community create and execute variants of published models and simulations to explore alternative simulation conditions, as well as help the community reuse published data visualizations to examine their results. We anticipate this platform will further bolster model reuse, composition, and collaboration.

### Additional modeling and simulation tools

Finally, we aim to help the community use runBioSimulations’ API to develop additional tools. For example, model repositories could use runBioSimulations to provide capabilities for executing their models, and model format developers could use runBioSimulations to implement test suites for verifying that simulators correctly support their formats.

## AVAILABILITY

The application and API are freely available without registration at https://run.biosimulations.org along with a tutorial, examples, and documentation. The source code is openly available under the MIT license at https://github.com/biosimulations/Biosimulations.

## ACKNOWLEDGEMENTS

We thank Arthur Goldberg, Herbert Sauro, James Schaff, Lucian Smith, and the Center for Reproducible Biomedical Modeling for input and feedback.

## FUNDING

This work was supported by National Institutes of Health award P41EB023912.

## Conflict of interest statement.

None declared.

## Notes

https://run.biosimulations.org

### Competing Interest Statement

The authors have declared no competing interest.

