## Supplementary Data for "runBioSimulations: an extensible web application that simulates a wide range of computational modeling frameworks, algorithms, and formats"

### SUPPORTED MODELING FRAMEWORKS, SIMULATION ALGORITHMS, MODEL FORMATS, AND SIMULATION TOOLS

Table S1 outlines the modeling frameworks, simulation algorithms, model formats, and simulation tools that runBioSimulations supports as of this writing (March 2021). Detailed, up-to-date information is available at https://biosimulators.org.

### ARCHITECTURE, IMPLEMENTATION, AND TESTING

#### Architecture

To execute a broad range of models, runBioSimulations has a modular, standards-driven architecture that abstracts the details of each modeling framework, simulation algorithm, and model format (Figure 3). The main components of runBioSimulations are back-end microservices for executing simulations on a high-performance computing (HPC) cluster, monitoring their progress, and collecting their results; a back-end database for saving simulation metadata and simulation results; an API which abstracts the microservices and database and provides external access to them; and a front-end GUI for submitting and monitoring simulations, and visualizing their results. Through the REST API, the front-end GUI functions as a client for the back-end microservices and back-end database. Importantly, the REST API is architected to support additional client applications. For example, investigators could use the API to programmatically execute simulations. Developers could use the API to create sophisticated third-party applications, such as an application that provides simulation capabilities to users of a model repository, a test suite similar to the SBML Test Suite (17) that verifies that simulators correctly support a model format, or a web-based application for constructing and executing models and simulations. Below, we describe the modular, standards-driven design of runBioSimulations through the workflow that it supports for running models.

First, the GUI displays a form for selecting a COMBINE archive (18) to simulate, selecting a simulation tool to execute the archive, selecting a version of that tool to utilize, and, optionally, entering an email address to receive notifications about the completion of the execution of the archive. The form retrieves the menu of available simulators from BioSimulators' REST API (https://biosimulators.org). Once the user submits the form, the GUI stores metadata about the simulation in the user's browser and uploads the COMBINE archive to runBioSimulations' REST API. The GUI uses this metadata to help users track their simulations.

After receiving a COMBINE archive, the REST API creates an entry for the simulation in the back-end database and instructs the simulation execution microservice to execute it. Next, the simulation execution microservice creates a high-performance computing (HPC) job to use the selected version of the selected simulator to run the uploaded COMBINE archive, and the microservice submits the job to the job manager for the HPC cluster. The simulation execution microservice can use all of the standardized simulators available from the BioSimulators registry because it listens to changes to BioSimulators; each time a standardized simulator is added or updated, the microservice pulls its Docker image and converts it to a Singularity image for execution on the HPC cluster.

While simulation jobs are running, the simulation monitoring microservice regularly polls the job manager to query the status of each running job and relays this status to the back-end database via the REST API. The GUI then polls this status from the

REST API and displays it in views for individual simulations and in a table that summarizes all of the simulations created by the user or shared with the user by other users.

When the monitoring microservice detects that a simulation job has completed, the microservice triggers the simulation results microservice to collect the data sets and execution log produced by the job, upload the results and log to the simulation database, and email the user about the completion of their simulation.

Once the data sets and log have been saved to the simulation database, the GUI can retrieve the data sets from the database for visualization, direct the user to download the data sets from the database, or retrieve the log from the database to render it for the user.

#### Implementation

We implemented the BioSimulators GUI in TypeScript, HTML, and SCSS using Angular, Material, and several additional packages. We used Ionic Storage to record the simulations available to each user (those created by the user or shared with them) within each user's browser. We used Plotly to visualize the results of simulations.

We implemented the simulation microservices in TypeScript using NATS, NestJS, SendGrid, Singularity, and SLURM. The execution microservice uses Singularity to pull Docker images for simulators and execute simulations, and the microservice uses SLURM to manage the execution of these simulations over an HPC cluster. The monitoring microservice regularly polls SLURM to detect when simulations start and terminate. The notification microservice uses SendGrid to email users about the completion of their simulations. We used NATS to coordinate the interactions between the microservices.

We implemented the simulation database in TypeScript using MongoDB, Amazon S3 object storage, Mongoose, and NestJS. The database uses MongoDB to store metadata about each simulation, such as the simulator which executed the simulation, when the simulation was submitted, and when it completed, as well as a log of the execution log of each simulation. The database also uses an Amazon S3-compatible object store to save the COMBINE archive for each simulation and the simulation results and other files produced by the execution of each archive. We used Mongoose and NestJS to define the schema for the MongoDB database and validate submissions to the database.

We implemented runBioSimulations' API in TypeScript using NestJS. We used Auth0 to manage access to the API.

We used Nx and GitHub to coordinate the development of the components of runBioSimulations.

#### Deployment

runBioSimulations is deployed using several cloud services and an HPC cluster and S3-compatible storage at the University of Connecticut School of Medicine (UConn; Figure 3). The GUI is deployed using Netlify. The simulations are executed on the HPC cluster, and all results and associated data are stored as objects in AWS-compatible S3 buckets. The simulation microservices and the REST API are deployed using a Kubernetes cluster run on Google Cloud. The simulation metadata database is deployed using MongoDB Atlas. Errors encountered during deployment are logged using Google Cloud Error Reporting.

#### Testing

We used ESLint to lint runBioSimulations, Jest to implement unit tests, and CodeCov to evaluate the coverage of our tests. To help us identify errors quickly, we used GitHub Actions to execute our tests each time we pushed changes to the runBioSimulations Git repository. We anticipate that this workflow will facilitate the continued development of runBioSimulations.

The simulation tools that runBioSimulations uses to execute simulations have been thoroughly tested by BioSimulators. Before accepting a simulator, BioSimulators verifies the simulator using a combination of automated testing and manual review. First, the BioSimulators test suite checks that the specifications of the simulators' capabilities (supported model formats, simulation algorithms, and modeling frameworks) is syntactically and semantically correct. Second, the BioSimulators test suite verifies that the simulator supports its reported capabilities, and that the simulator runs simulations and generates reports of their results consistently with BioSimulators' conventions. This is achieved by using the simulator to execute a collection of curated and synthetically generated COMBINE archives that probe support for features of the COMBINE archive format, SED-ML, and the model languages that the simulator supports, such as the BioNetGen Language (BNGL, 1) and the Systems Biology Markup Language (SBML, 4). Third, one of BioSimulators' curators manually reviews the specifications of the simulator and the results of the automated tests. Furthermore, most of the simulators used by runBioSimulations have been verified using the SBML Test Suite (19).

### CASE STUDY: ASSESSING THE PRACTICAL REUSABILITY OF PUBLISHED SIMULATION PROJECTS TO INDIVIDUAL INVESTIGATORS

In this section, we present two case studies that illustrate the utility of runBioSimulations' ability to execute a broad range of simulations.

Because runBioSimulations has unique capabilities to execute a broad range of simulation studies and to collect structured logs that facilitate the debugging of complex simulations, runBioSimulations is uniquely positioned for evaluating the reproducibility and, in turn, reusability and composability of existing simulation projects. We used runBioSimulations to assess the reproducibility of simulation projects published through BioModels (20), one of the largest collections simulation projects, to individual investigators. In contrast, most previous studies have focused on the reproducibility of published simulation projects to expert curators, who have experience and the time to debug problems with simulations published by other investigators.

First, we used BioModels' REST API to determine which BioModels entries include simulation experiments. As of the time we searched BioModels (Spring 2020), we determined that 214 BioModels entries include simulation experiments in SED-ML format. Second, we used the REST API to download these projects in the COMBINE archive format. Third, we used our simulation capabilities to execute these projects.

Next, we examined the results of the execution of these COMBINE archives to evaluate their ability to reproduce published simulation results. This revealed that only two ($<1\%$) of these simulation projects are executable.

Next, we collected the execution logs of these simulation projects to dissect the proximate reasons why most of these COMBINE archives cannot be successfully executed. We found that 84% of the SED-ML files in these COMBINE archives have broken references to model files (the model in the COMBINE archive is located at a different path than that indicated in the SED-ML file); that 15% of the SED-ML files in these archives are syntactically invalid, most commonly due to lacking required attributes, having duplicate values of attributes which must have unique values, and having attributes with invalid values; that BioModels' API cannot generate COMBINE archives for 0.5% of entries; and that 0.5% of the projects cannot be executed because their SED-ML files include broken references to variables of models (invalid XPATHs for targets of SED-ML variables). We also examined the results of the two COMBINE archives that did successfully execute to gain insights into best practices for publishing reproducible simulation projects. This revealed another issue, that the simulation experiments in one of the COMBINE archives referenced models outside the archive. While this makes that COMBINE archive executable, this design is inconsistent with BioModels' goal of exporting COMBINE archives that standalone, containing simulations and all of the models required for their execution.

Next, we used BioSimulations utils, one of the libraries we developed to implement runBioSimulations, to dissect these COMBINE archives and identify additional issues beyond those identified above that would prevent the successful execution of these archives. This revealed additional issues with 20% of the simulation experiments contained in BioModels. Many of these issues fall into the same categories outlined above. In addition, some of the simulation experiments have invalid entity references (e.g., references to undefined models) and semantically-ambiguous axes of plots.

Next, we used BioSimulations utils to automatically clean the issues outlined above. This enabled us to produce syntactically and semantically-valid simulation experiments for 97% of entries in BioModels. The remaining 3% of archives have more complex problems that are more difficult to clean programmatically.

Next, we executed these cleaned simulation experiments. This revealed that 10% of the simulation experiments could not be executed with additional simulation tools beyond the original tools that were used to create the simulation experiments.

Other recent studies have also found that many published simulation experiments are not reproducible. For example, Tiwari et al. found that almost 50% of published simulation experiments provide enough information for their reproduction (21). However, Tiwari et al. needed to expend significant effort to reproduce many simulation experiments, debugging experiments, sometimes with assistance from their original authors. As a result, many previous studies have estimated the reproducibility of published simulation experiments to expert curators rather than individual investigators. In practice, reusing simulations requires investigators to be able to quickly execute simulation experiments with modest effort. BioModels and other repositories aim to facilitate this by providing verified simulation experiments in community formats such as COMBINE and SED-ML. Our study mimicked this scenario by focusing on curated files that are readily available to investigators, without additional debugging in previous studies by expert curators. As a result, we believe our findings better reflect the reusability of published simulation experiments to individual investigators.

Going forward, we aim to help the developers of BioModels and simulation tools address the issues identified above.

### COMPARISON WITH OTHER ONLINE SIMULATION TOOLS

Numerous software tools help investigators execute simulations, visualize their results, and share these resources (Table S2). Most of these tools focus on one or two model formats, one or two modeling frameworks, and one or a few simulation algorithms. In contrast, runBioSimulations provides a single, consistent, extensible portal for executing simulations that involve multiple formats, frameworks, and simulation algorithms.

This is achieved through a novel standards-driven architecture. In particular, runBioSimulations is one of only a few tools that can execute SED-ML files and the first tool that leverages BioSimulators. This focus on community resources makes the simulation logic of runBioSimulations transparent and transferable beyond runBioSimulations, and it makes runBioSimulations extensible to community contributions of additional modeling frameworks, simulation algorithms, and model formats.

runBioSimulations is also one of only a few web-based simulation tools that can store simulation results. We believe this makes runBioSimulations well-suited for sharing computationally-expensive simulations.

Finally, runBioSimulations is one of only a few tools that enable simulations and their results to be shared via URLs similar to cloud storage systems such as Google Drive. We believe this simple sharing mechanism is easy for users to adopt.

### COMMUNITY FEEDBACK, INPUT, AND CONTRIBUTIONS

We welcome community feedback, input, and direct contributions to runBioSimulations. A guide to contributing to runBioSimulations is available at https://github.com/biosimulations/Biosimulations. We also encourage developers to use the runBioSimulations API to develop additional applications. For example, developers could use the API to develop web-based tools for building and simulating models or build test suites for verifying that simulators correctly support model languages.

BioSimulators also welcomes the community to contribute additional simulators for additional modeling frameworks, simulation algorithms, and model formats. Information about submitting simulators is available at https://biosimulators.org.

**Table S1. Overview of the modeling frameworks, simulation algorithms, model formats, and simulation tools supported by runBioSimulations as of March 2021.** Detailed, up-to-date information is available at https://biosimulators.org.

| Model format | | Modeling framework | Simulation algorithm | Simulator | |
|---|---|---|---|---|---|
| BNGL | 1 | continuous kinetic ODE | CVODE | BioNetGen | 2 |
| BNGL | 1 | discrete kinetic | Gillespie direct algorithm | BioNetGen | 2 |
| BNGL | 1 | discrete kinetic | Network-free simulation method | BioNetGen/NFSim | 2, 3 |
| BNGL | 1 | discrete kinetic | Partitioned leaping method | BioNetGen | 2 |
| SBML | 4 | continuous kinetic DAE | IDA | VCell | 5, 6 |
| SBML | 4 | continuous kinetic ODE | Adams-Moulton method | VCell | 5, 6 |
| SBML | 4 | continuous kinetic ODE | CVODE | PySceS | 7 |
| SBML | 4 | continuous kinetic ODE | CVODE | tellurium | 8 |
| SBML | 4 | continuous kinetic ODE | CVODE | VCell | 5, 6 |
| SBML | 4 | continuous kinetic ODE | CVODES | AMICI | 9 |
| SBML | 4 | continuous kinetic ODE | Dormand-Prince 8(5,3) method | GillesPy2 | 10 |
| SBML | 4 | continuous kinetic ODE | Dormand-Prince 4(5) method | GillesPy2 | 10 |
| SBML | 4 | continuous kinetic ODE | Euler forward method | VCell | 5, 6 |
| SBML | 4 | continuous kinetic ODE | Fehlberg method | tellurium | 8 |
| SBML | 4 | continuous kinetic ODE | Fehlberg method | VCell | 5, 6 |
| SBML | 4 | continuous kinetic ODE | Fourth-order Runge-Kutta method | tellurium | 8 |
| SBML | 4 | continuous kinetic ODE | Fourth-order Runge-Kutta method | VCell | 5, 6 |
| SBML | 4 | continuous kinetic ODE | LSODA | GillesPy2 | 10 |
| SBML | 4 | continuous kinetic ODE | LSODA | PySceS | 7 |
| SBML | 4 | continuous kinetic ODE | LSODA/LSODAR | COPASI | 11 |
| SBML | 4 | continuous kinetic ODE | Midpoint method | VCell | 5, 6 |
| SBML | 4 | continuous kinetic ODE | Milstein method | VCell | 5, 6 |
| SBML | 4 | continuous kinetic ODE | Newton-type method | tellurium | 8 |
| SBML | 4 | continuous kinetic ODE | Radau method | COPASI | 11 |
| SBML | 4 | continuous kinetic ODE | VODE | GillesPy2 | 10 |
| SBML | 4 | continuous kinetic ODE | ZVODE | GillesPy2 | 10 |
| SBML | 4 | discrete kinetic | Gibson-Bruck Next Reaction Method | COPASI | 11 |
| SBML | 4 | discrete kinetic | Gibson-Bruck Next Reaction Method | VCell | 5, 6 |
| SBML | 4 | discrete kinetic | Gillespie direct algorithm | COPASI | 11 |
| SBML | 4 | discrete kinetic | Gillespie direct algorithm | GillesPy2 | 10 |
| SBML | 4 | discrete kinetic | Gillespie direct algorithm | tellurium | 8 |
| SBML | 4 | discrete kinetic | Adaptive explicit-implicit tau-leaping method | COPASI | 11 |
| SBML | 4 | discrete kinetic | tau-leaping method | COPASI | 11 |
| SBML | 4 | discrete kinetic | tau-leaping method | GillesPy2 | 10 |
| SBML | 4 | hybrid kinetic | Adaptive Gibson/Milstein Method | VCell | 5, 6 |
| SBML | 4 | hybrid kinetic | Gibson/Euler-Maruyama Method | VCell | 5, 6 |
| SBML | 4 | hybrid kinetic | Gibson/Milstein Method | VCell | 5, 6 |
| SBML | 4 | hybrid kinetic | Hybrid tau-leaping Method | GillesPy2 | 10 |
| SBML | 4 | hybrid kinetic | Next Reaction method/LSODA method | COPASI | 11 |
| SBML | 4 | hybrid kinetic | Next Reaction method/RK-45 method | COPASI | 11 |
| SBML | 4 | hybrid kinetic | Next Reaction method/Runge-Kutta method | COPASI | 11 |
| SBML | 4 | stochastic kinetic SDE | Second order Runge-Kutta method | COPASI | 11 |
| SBML-fbc | 12 | flux balance | Flux balance analysis | CBMPy | 13 |
| SBML-fbc | 12 | flux balance | Flux balance analysis | COBRApy | 14 |
| SBML-fbc | 12 | flux balance | Flux variability analysis | CBMPy | 13 |
| SBML-fbc | 12 | flux balance | Flux variability analysis | COBRApy | 14 |
| SBML-fbc | 12 | flux balance | Geometric flux balance analysis | COBRApy | 14 |
| SBML-fbc | 12 | flux balance | Parsimonious FBA (minimum sum of fluxes) | CBMPy | 13 |
| SBML-fbc | 12 | flux balance | Parsimonious FBA (minimum sum of fluxes) | COBRApy | 14 |
| SBML-fbc | 12 | flux balance | Parsimonious FBA (minimum number of active fluxes) | CBMPy | 13 |
| SBML-qual | 15 | logical | Asynchronous logical model simulation method | BoolNet | 16 |
| SBML-qual | 15 | logical | Probabilistic logical model simulation method | BoolNet | 16 |
| SBML-qual | 15 | logical | Synchronous logical model simulation method | BoolNet | 16 |
| SBML-spatial | | spatial discrete kinetic | Brownian diffusion Smoluchowski method | VCell | 5, 6 |
| SBML-spatial | | spatial continuous kinetic PDE | Fully-Implicit Finite Volume | VCell | 5, 6 |
| SBML-spatial | | spatial continuous kinetic PDE | Semi-Implicit Finite Volume-Particle Method | VCell | 5, 6 |

**Table S2. Comparison of the features of runBioSimulations and select other online simulation tools.** The ‘Mathematical frameworks,’ ‘Formalisms,’ and ‘Model formats’ columns indicate the number of simulation algorithms supported by each tool. The ‘Other formats‘ columns indicate which tools support SED-ML and the COMBINE archive format. The ‘Collaborative features’ columns indicate which tools help users build and share simulations and their results. Question marks indicate information that we could not obtain from public documentation. Filled checkmarks indicate fully supported features; open checkmarks indicate partially supported features; circles indicate algorithms that the community could contribute to runBioSimulations.

| | | Mathematical frameworks | | | | | | | | Form-alisms | | Model formats | | | | | | | Other formats | | Collaborative features | | | | | | | | |
|---|---|---|---|---|---|---|---|---|---|---|---|---|---|---|---|---|---|---|---|---|---|---|---|---|---|---|---|---|---|
| Tool | Ref | *Kinetic, DAE* | *Kinetic, ODE/PDE* | *Kinetic, SDE* | *Kinetic, Discrete* | *Kinetic, Hybrid* | *Kinetic, Steady-state* | *Flux-balance* | *Logical* | *Spatial* | *Rule-based* | *BNGL (1)* | *Kappa (22)* | *NeuroML (23)* | *SBML (4)* | *SBML-fbc (12)* | *SBML-qual (15)* | *SBML-spatial* | *SED-ML (24)* | *COMBINE (18)* | *Build models* | *Build simulations* | *Store simulations* | *Store results* | *Sharing with users* | *Sharing via URLs* | *Publication* | *Modular, local simulation* | *Extensible to community* |
| BioUML | 25 | | | | | | | 4 | 2 | | | | | | | 4 | 2 | | | | | ✔ | ✔ | ✔ | ✓ | | ✓ | ✓ | ✓ |
| Escher-FBA | 26 | | | | | | | 1 | | | | | | | | 1 | | | | | | ✔ | | | | | | | |
| Fluxer | 27 | | | | | | | 1 | | | | | | | 1 | | | | | | | | | | | | | | |
| MetaNetX | 28 | | | | | | | 3 | | | | | | | 3 | | | | | | | | | | | | | | |
| JWS Online | 29 | | 1 | | | | 1 | 1 | | | | | | | 2 | 1 | | | ✓ | ✓ | ✔ | ✔ | ✓ | | | | ✓ | | |
| KappaTools | 30 | | | | 1 | | | | | | | | 1 | | | | | | | | ✔ | ✔ | | | | | | ✔ | |
| KBase | 31 | | | | | | | 1 | | | | | | | | 1 | | | | | | | ✔ | ✔ | ✔ | | ✔ | | |
| Open Source Brain | 32 | | ? | ? | | ? | | | | ? | | | | ? | | | | | | | | ✔ | ✔ | | ✓ | | ✔ | ✓ | ✓ |
| runBioSimulations | | 1 | 15 | 1 | 5 | 7 | 1 | 5 | 3 | 6 | 4 | 4 | ○ | ○ | 23 | 5 | 3 | 6 | ✔ | ✔ | | | ✔ | ✔ | | ✔ | | ✔ | ✔ |
| StochSS | 10 | | 1 | | 2 | 1 | | | | | | | | | 4 | | | | | | ✔ | ✔ | ✔ | ✔ | | | | ✓ | |
| The Cell Collective | 33 | | | | | | | 4 | 2 | | | | | | | 4 | 2 | | | | ✔ | ✔ | ✔ | | ✔ | ✔ | ✔ | | |
